# Multiple long-range *cis* interactions generate CTCF insulator-dependent viral chromatin domains in quiescent HSV-1 genomes

**DOI:** 10.1101/2025.05.08.652945

**Authors:** Alyssa Richman, Sophie Kogut, Terri Edwards, Joseph Boyd, Princess Rodriguez, Michael Mariani, Mason A. Shipley, Kayley A. Manuel, Ziyun A. Ye, David C. Bloom, Seth Frietze, Donna M. Neumann

## Abstract

In cellular genomes, CTCF insulators impact transcription over small distances in a one-dimensional manner and over much longer distances in a three-dimensional manner by maintaining chromatin loops. We have previously shown that the latent HSV-1 genome contains CTCF insulators that function to regulate lytic transcription of adjacent genes in a one-dimensional manner. Here, we test the hypothesis that HSV-1 CTCF insulators nucleate chromatin loops to regulate the expression of distance separated gene regions through three-dimensional organization of viral genomes. We used 4C-*seq* methods to identify multiple long-range *cis* interactions in HSV-1 genomes that generate viral chromatin domains, including those nucleated by the viral CTCF insulator CTRL2. Deletion of the CTRL2 insulator disrupted these viral chromatin domains. Loop-nucleating interactions were quantitated with a novel approach (UMI-4C-*seq*) that utilizes unique molecular identifiers to label and count chromatin interactions associated with specific viewpoint primers. *Cis*-interaction peaks across four different viewpoints were quantified. Viral genomes lacking CTRL2 displayed more *cis*-interaction peaks and wider ranges of interaction lengths compared to wt virus, suggesting altered chromatin organization. Furthermore, differential looping analysis showed viral genomes lacking CTRL2 displayed a more transcriptionally permissive chromatin environment. Thus, the CTRL2 insulator functions as a critical regulator of long-range chromatin interactions and its deletion reshapes the viral chromatin landscape, leading to a more accessible and dynamic regulatory environment that may influence HSV-1 transcriptional programs and latency-associated chromatin states.

## Introduction

Herpes Simplex Virus 1 (HSV-1) establishes a lifelong infection in ∼70% of adults and is a significant human pathogen with clinical manifestations that range from herpes labialis to keratitis, and in rare cases, death from HSV-1 induced encephalitis [1–3]. Following the primary lytic infection, HSV-1 establishes latency in sensory neurons, where the viral genome is essentially silenced. HSV-1 periodically reactivates from latent reservoirs in response to various physiologic and environmental stressors, and repeated reactivation can result in ocular pathogenesis and corneal blindness over time [2–4]. Currently, the therapeutics available to treat HSV-1 infections are limited to antivirals that target replicating virus, leaving the latent viral reservoirs untouched, largely because the mechanisms that govern the establishment, maintenance, and exit from latency remain under-defined.

HSV-1 genomes are organized into distinct chromatin domains where the immediate early (IE) lytic genes are maintained in transcriptionally repressed domains enriched in the heterochromatic H3K27me3 and H3K9me3 histone markers [5–12], while the non-coding RNA LAT (Latency Associated Transcript) is enriched in euchromatin H3K9K14 and H3K4me2/3 during latency [5, 13–20]. The segregation and maintenance of these distinctly different latent chromatin domains are maintained by functional insulator elements commonly known as CTCF insulators [21, 22]. CTCF insulators consist of a conserved DNA binding motif to which CTCF proteins bind to elicit insulator function. In eukaryotic cells, CTCF insulators are considered “master regulators” of transcription [23, 24], controlling gene expression through a myriad of mechanisms, some of which include acting as enhancer-blockers that prevent inappropriate promoter activation or as barrier elements that prevent heterochromatin spread [23–29]. Using an algorithm against the HSV-1 sequence that recognized a conserved reiterated DNA motif that CTCF proteins bind to (CCCTC/CTCCC sequence motifs), we identified and then subsequently characterized seven CTCF insulators in latent HSV-1 genomes [21, 22, 30, 31]. Interestingly, 6 of the 7 CTCF insulators identified were in the repeat regions of HSV-1 and strikingly, they flanked not only the 5’ exon region of LAT, a region that contains the LAT promoter and enhancer elements that are required for efficient reactivation [32], but also the IE genes [21], suggesting that these insulators were key elements in maintaining gene silencing during the latent infection.

CTCF insulators also dimerize to form three-dimensional (3D) structures known as chromatin loops. These long-range *cis* interactions bring distance separated enhancers and promoters together in close spatial proximity for transcriptional control [27, 33–35]. Chromatin loops are found in eukaryotic cells and beta and gamma herpesviruses. Further, these 3D loop structures are important for transcriptional control, as they regulate latency types in EBV or change/rearrange in response to reactivation in KSHV [36–40]. Chromatin loops are anchored and stabilized by the cohesin protein complex [39, 41–44], and our recent findings that showed HSV-1 encoded CTCF insulators also colocalized with cohesin complex proteins suggested that HSV-1 genomes were also organized into 3D chromatin loops during latency [45]. To determine if these 3D chromatin structures were in HSV-1 genomes, we leveraged well-defined circular chromosome conformation capture assays combined with sequencing (4C-*seq*) that have been previously used to show chromatin loop organization of other DNA viruses [38, 46–50].

Using HSV-1 Lund human mesencephalic (LUHMES) neuronal cells quiescently infected with wild-type (wt) HSV-1 strain 17Syn+, we identified multiple long-range *cis* interactions by 4C-*seq* methods [47, 48] in HSV-1 genomes that generate viral chromatin domains. To determine whether the 3D chromatin structure of the wt genome was dependent on individual virally encoded insulators, we leverage our well characterized recombinant virus containing a small 130bp deletion of the core binding domain of the CTRL2 insulator, a functional insulator downstream from the LAT enhancer element (ΔCTRL2) [31, 45, 51, 52]. In LUHMES quiescently infected with the ΔCTRL2 recombinant, we showed that deletion of the CTRL2 insulator of HSV-1 resulted in the loss of a specific long-range *cis* interaction that mapped to the US region of the viral genome near the US8 and US9 overlapping genes, two genes that are required for efficient anterograde axonal transport in reactivation. Taken together, these results suggest that the 3D chromatin structure of the latent viral genome is important for the virus’s ability maintain latency and to reactivate.

To further quantify both the abundance of and the changes in long-range interactions observed between the wt and ΔCTRL2 genomes, we then optimized UMI-4C-*seq* (Unique Molecular Identifier 4C-sequencing) in our LUHMES model. UMI-4C-*seq* is a technique that provides a robust method of capturing and quantifying long-range interactions compared to traditional 4C-*seq* by incorporating unique molecular identifiers (UMIs) that improve the precision and accuracy of detecting interactions in complex populations of cells. We quantitated changes in long-range interactions of the ΔCTRL2 virus compared to wt using viewpoint primers proximal to VP16, LAT, CTRL2, and ICP4 loci by UMI-4C. We showed that the deletion of the CTRL2 insulator resulted in significant and quantifiable changes to the 3D chromatin structure of viral genomes, where numerous novel interactions were gained and lost in the ΔCTRL2 recombinant. These findings reflect a disruption of local chromatin organization that is observed in wt genomes during HSV-1 quiescence. Combined, our data suggests that the 3D structure of HSV-1 genomes are important for regulated gene expression during latency and for organizing HSV-1 in a structural conformation that is favorable for reactivation from latency.

## Results

### 4C-seq analysis of HSV-1 identifies *cis* chromatin contacts in quiescently infected human neuronal cells

To elucidate the chromatin architecture of HSV-1 genomes in neurons quiescently infected with HSV-1, we performed 4C-*seq* experiments using human LUHMES cells infected with 17Syn+ [53, 54]. 4C-*seq* viewpoint primer bait sequences were designed to detect the interaction frequencies between single viewpoint DNA fragments and *cis*-interactions with distal regions of the HSV-1 genome. We selected distinct 4C-*seq* viewpoints that mapped to either the unique long (UL) or the unique short (US) regions of the HSV-1 genome (located at 28 kb and 133 kb, respectively) (**Fig. 1**). The baits targeting viewpoints 1 and 2 (VP1 and VP2, **Fig. 1A and 1B**) are proximal to the immediate–early (IE) genes US1, encoding the ICP22 repressor, and the UL13 early gene encoding the protein kinase. RNA-*seq* analysis of HSV-1 infected cells has demonstrated that both ICP22 and UL13 are expressed during the replication cycle and are involved in viral gene expression and replication competence [55]. The viewpoints were specific to HSV-1 infected cells and failed to amplify non-infected 4C templates (**S. Fig.1**). Three independent 4C-*seq* experiments were conducted and sequenced to a depth of >1 million reads per 4C library. Peak calling analysis was performed to identify significant *cis*-chromatin contacts for each viewpoint (see methods). We identified 11 significant peaks (FDR < 0.1) corresponding to VP1 *cis*-contacts that spanned the length of the HSV-1 genome, including two clusters of regions at 80-90 and 130-140 kb distal to the VP1 region of US1 (**Fig. 1A**). We similarly identified 10 peaks with VP2 that showed clustering within the 80-130 kb region (**Fig. 1B**). Notably, the *cis*-contacts for both viewpoints showed a high degree of concordance (**Fig. 1C**), where >85% of peaks were shared between viewpoint baits, suggesting central chromatin domains are formed by HSV-1 **(S. Table 1**).

**Figure 1:**
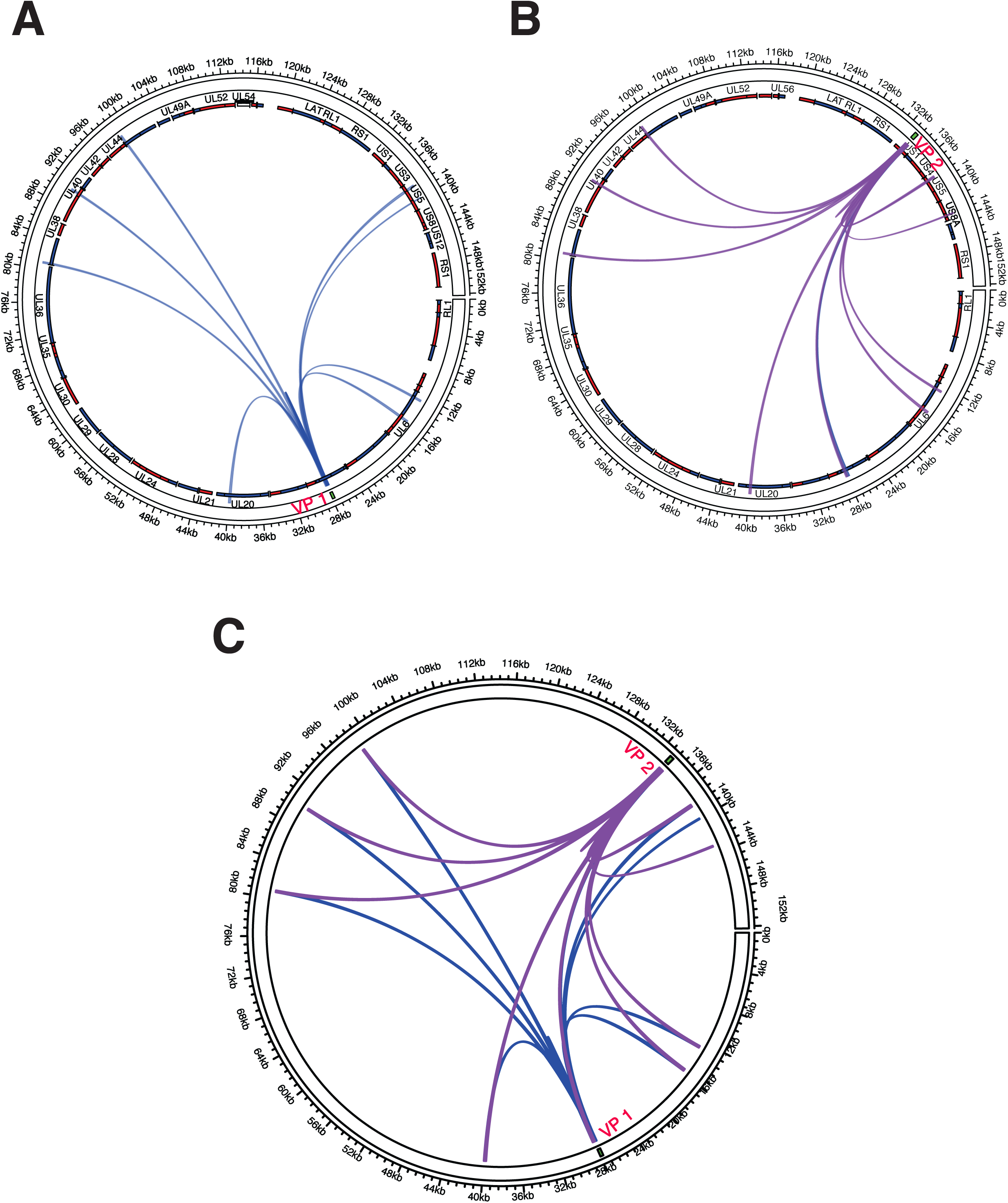
4C-*seq* analysis of *cis*-chromatin contacts within the HSV-1 genome. 4C-*seq* viewpoint primers (VP1 and VP2) targeting genomic regions on the circularized 152 kb HSV-1 genome. **A.** Circos plots showing viral open reading frames (ORFs) in sense (red) and antisense (blue) strands, significant *cis*-interactions (peaks) identified with VP1 bait regions. **B.** Circos plots showing HSV-1 viral open reading frames (ORFs) in sense (red) and antisense (blue) strands, significant *cis*-interactions (peaks) identified with VP2 bait regions. **C.** Merged plots of both VP1 and VP2 in wild-type (wt-17Syn+) genomes.

### Long-range interactions in HSV-1 shift in the absence of ΔCTRL2

CTCF and associated insulator elements regulate the formation of transcriptional chromatin domains. To determine potential changes in HSV-1 chromatin structure associated with the deletion of a viral CTCF insulator element, we leveraged our recombinant ΔCTRL2 virus, a mutant virus containing a 130bp deletion of the core CTCF binding site known as CTRL2 downstream from the LAT 5’exon [51]. We had previously shown that the CTRL2 insulator was required to maintain IE gene silencing during latency through the maintenance of heterochromatin on the HSV-1 genome [31, 51], suggesting that the viral insulator element was integral to the regulation of latent HSV-1 chromatin domains. Using 4C-*seq* on LUHMES cells quiescently infected with ΔCTRL2, we identified 12 and 14 significant peaks for VP1 and VP2 respectively with ΔCTRL2 (**Fig. 2**). The interactions for VP1 and VP2 both showed near complete overlap with those identified in LUHMES cells infected with wt virus. However, two additional *cis*-interactions with the distal region of the HSV-1 genome were identified with VP1 in ΔCTRL2 compared to wt samples **(Fig. 2A, S**. **Table 1**). Notably, VP1 interacted with a domain spanning 143-152kb in ΔCTRL2 which was absent in wt virus (**Fig. 2B)**. In addition, one specific interaction was lost in ΔCTRL2 compared to wt virus (VP2 and 143kb, corresponding to US8 and US9 overlapping regions **Fig. 2-see arrow).** These 4C-*seq* results indicate that HSV-1 transcriptional domains may shift in their composition in latent infections when insulator domains are deleted in a manner that would impact the ability to reactivate.

**Figure 2:**
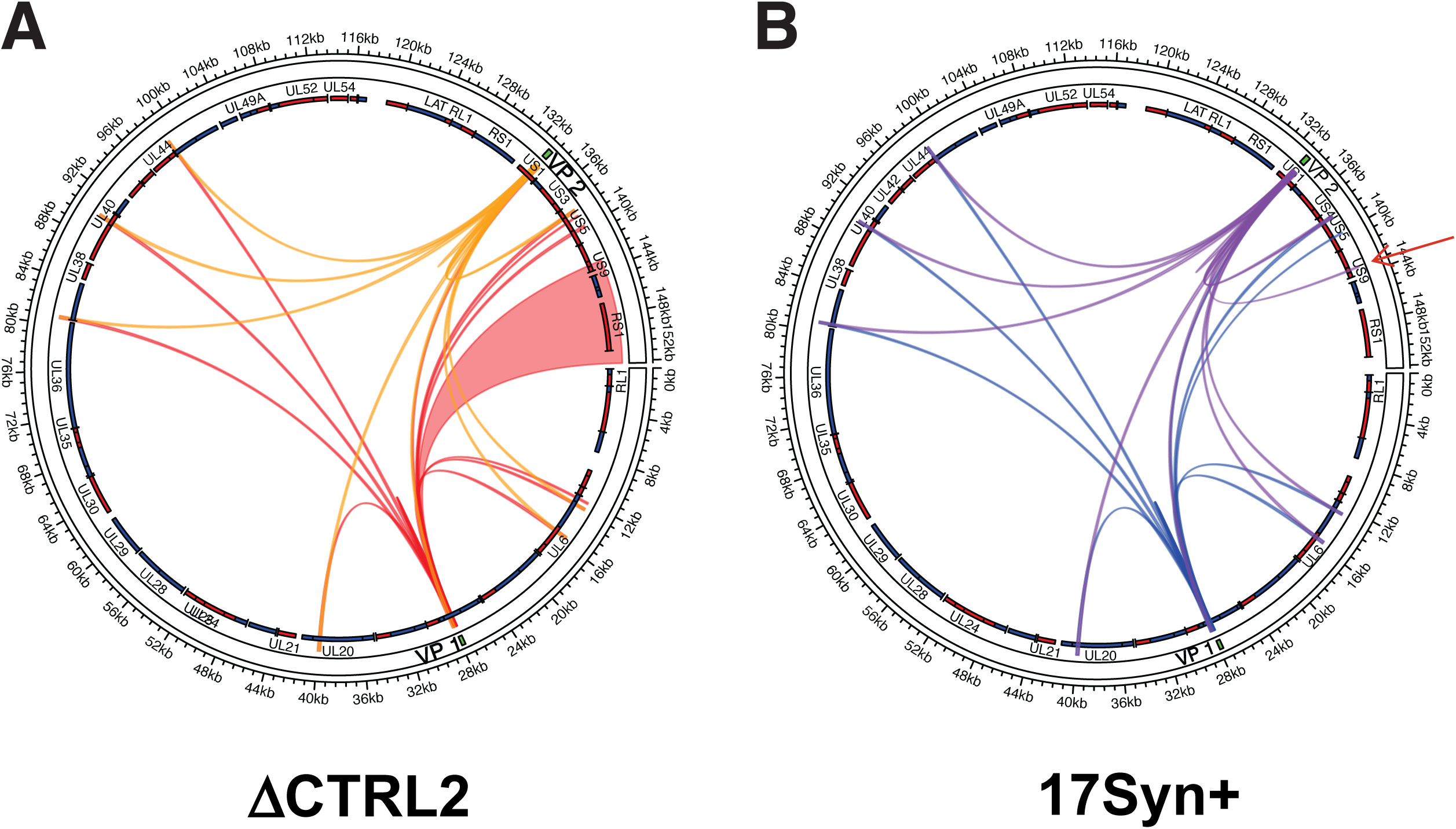
*Cis*-chromatin contacts of HSV-1 ΔCTRL2 strain lacking the functional viral CTRL2 insulator. **A.** 4C-*seq* analysis using both viewpoint primers VP1 and VP2 comparing LUHMES cells quiescently infected with the HSV-1 ΔCTRL2 virus strain harboring a targeted deletion in the CTCF insulator domain. The area shaded in red indicates interactions gained in the recombinant virus compared to wt virus (shown in panel B). **B.** The wild-type (wt) 17Syn+ HSV-1 strain is shown for comparison. Arrow points to the interaction that is lost upon deletion of the viral CTRL2 insulator. This interaction maps to anterograde transport genes US* and the overlapping US( region of the HSV-1 genome. Shown in both graphs are all significant peaks for both VP1 and VP2 bait regions.

### UMI-4C-*seq* identified differential chromatin interactions in HSV-1 ΔCTRL2 compared to wt virus

To further analyze the changes in chromatin loops between wt and ΔCTRL2 viruses, we performed UMI-4C experiments. This technique combines chromosome conformation capture with unique molecular identifiers (UMI) to quantitatively compare the differential analysis of targeted HSV-1 *cis*-contact profiles [56]. We designed primers targeting distinct HSV-1 bait regions proximal to the viral genes/loci including CTRL2, ICP4, LAT, and VP16. UMI-4C libraries were generated using independent LUHMES cultures quiescently infected with wt or ΔCTRL2. Viewpoint primers targeting distinct regions of the HSV-1 genome proximal to VP16, LAT, CTRL2, and ICP4 loci are shown in **S. Table 2**. Paired-end sequencing of multiplexed UMI-4C amplicons resulted in a similar number of reads with a relatively balanced distribution of UMIs and a high degree of consistency across replicates across each bait (**S. Fig. 3A**). UMI-4C analysis identified *cis*-interaction peaks across viewpoints with differences in frequency and length between wt and ΔCTRL2 genomes (**Fig. 3A&B)**. For the VP16 viewpoint, the results show several new contacts in the ΔCTRL2 samples that are absent in the wt genomes, particularly in regions spanning 90-110 kb and 130-150 kb (**Fig. 3F)**. This demonstrates that the deletion of the CTCF insulator element has a substantial impact on the chromatin interactions around the VP16 region. In the ΔCTRL2 samples, there is also an increase in contact frequencies at regions proximal to LAT, suggesting potential changes in chromatin architecture linked to the deletion of the CTCF insulator **(S. Fig. 3B&C)**. The trend plot highlights these differential contact intensities, showing peaks that indicate higher interaction frequencies in the ΔCTRL2 samples compared to wt **(Fig. 3C&D)**. Overall, ΔCTRL2 genomes exhibited more *cis*-interaction peaks at most viewpoints compared to wt (**Fig. 3A**). For instance, the ΔCTRL2 virus showed nearly twice as many significant interactions at LAT (19 vs. 10) and VP16 (7 vs. 3), whereas wt exhibited a higher number of peaks at CTRL2 (11 vs. 8). ΔCTRL2 interactions were generally longer on average across viewpoints, with notable differences at CTRL2 (3474 bp vs. 2101 bp) and LAT (3771 bp vs. 2744 bp) (**Fig. 3B**). The range of interaction lengths was consistently broader in ΔCTRL2 genomes, suggesting altered chromatin organization compared to wt genomes **(S. Table 3)**.

**Figure 3.**
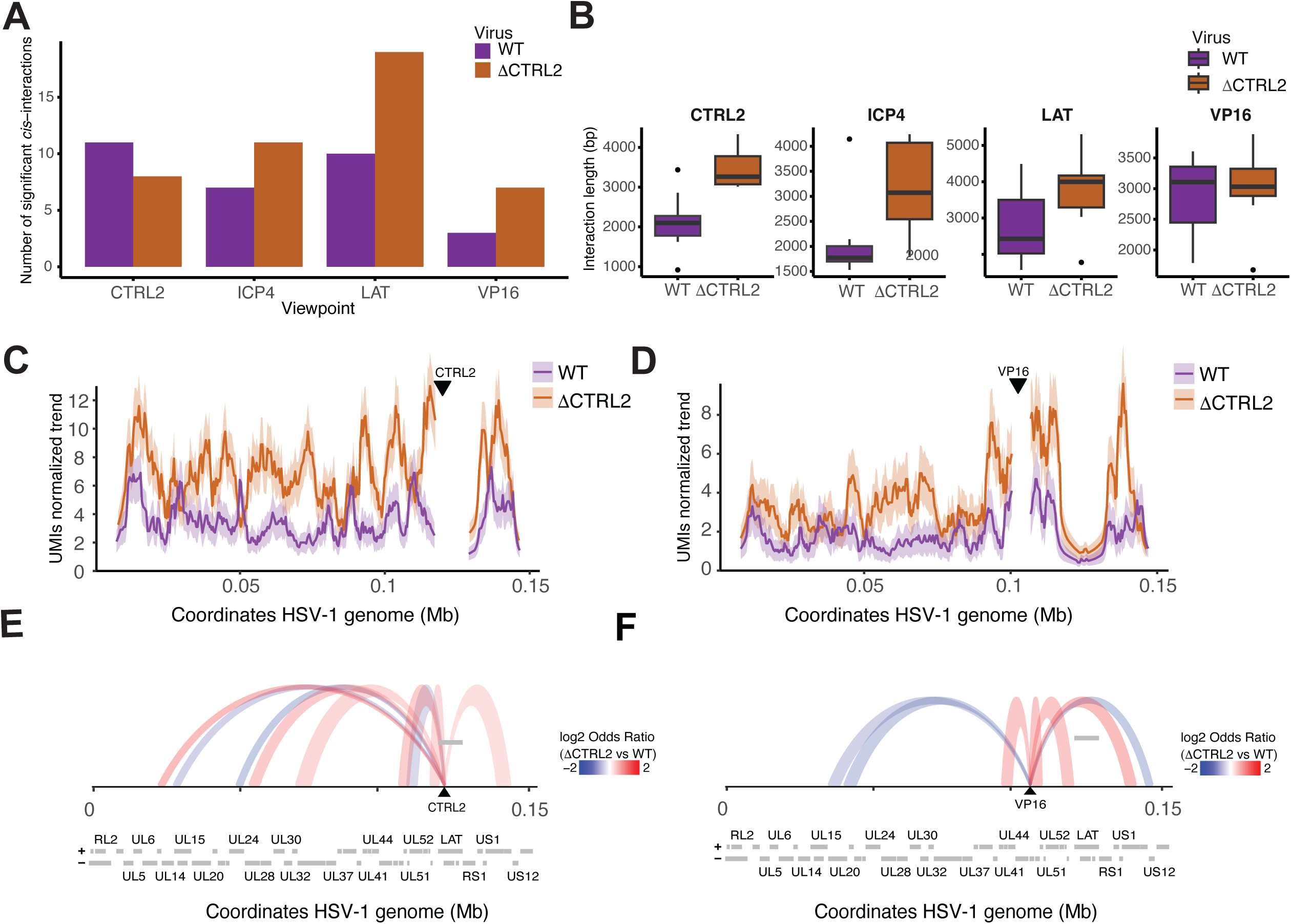
UMI-4C-*seq* analysis reveals altered *cis*-chromatin architecture in latent HSV-1 genomes upon deletion of the CTRL2 insulator. **A**. Number of significant *cis*-interactions (FDR < 0.1) identified at each UMI-4C viewpoint (CTRL2, ICP4, LAT, and VP16) in wild-type (wt; purple) and CTRL2-deleted (ΔCTRL2; orange) HSV-1 genomes. ΔCTRL2 samples show an overall increase in the number of long-range interactions, particularly at LAT and VP16 viewpoints. **B**. Distribution of interaction lengths (in base pairs) for each viewpoint in wt and ΔCTRL2. **C.** UMI-normalized interaction frequency profiles across the HSV-1 genome for CTRL2. **D.** UMI-normalized interaction frequency profiles across the HSV-1 genome VP16 (right) viewpoints in wt and ΔCTRL2-infected LUHMES cells. Increased interaction frequencies and broader interaction domains are observed in ΔCTRL2 genomes relative to wt. **E.** Differential chromatin interaction arcs for CTRL2 viewpoints visualized as arc plots across the HSV-1 genome. Arcs represent statistically significant changes in *cis*-interactions peaks between ΔCTRL2 and wt genomes. Red arcs indicate **increased** interaction frequencies in ΔCTRL2 relative to wt (log odds ratio > 1), while blue arcs indicate **reduced** interactions. HSV-1 genome coordinates and annotated ORFs are displayed below the arc plots. **F.** Differential chromatin interaction arcs VP16 viewpoints visualized as arc plots across the HSV-1 genome. Arcs represent statistically significant changes in *cis*-interactions peaks between ΔCTRL2 and wt genomes. Red arcs indicate increased interaction frequencies in ΔCTRL2 relative to wt (log odds ratio > 1), while blue arcs indicate reduced interactions. HSV-1 genome coordinates and annotated ORFs are displayed below the arc plots.

Differential looping analysis of UMI-4C data comparing the ΔCTRL2 recombinant to wt 17Syn+ genomes showed ΔCTRL2 had significantly altered chromatin interactions across analyzed HSV-1 loci, including VP16 and CTRL2 (**Fig. 3E&F**). At the VP16 locus, ΔCTRL2 genomes exhibited increased distal interactions and a broader *cis*-interactions compared to wt genomes, consistent with the loss of boundary constraints imposed by the CTRL2 element. Similarly, at the proximal CTRL2 locus, quiescent ΔCTRL2 genomes displayed an expansion of interaction domains and enhanced interaction frequencies flanking the locus, indicative of a more permissive chromatin environment. These changes likely reflect a disruption of local chromatin organization, potentially facilitating greater transcriptional accessibility and regulatory interplay. Differential analysis across all loci showed that ΔCTRL2 genomes consistently exhibited more extensive and long-range interactions than wt, supporting the insulator role in preserving chromatin compartmentalization and stability (**Fig 3E&F)**. The *cis*-chromatin interactions at LAT and ICP4 loci also demonstrated genome-wide effects of CTRL2 deletion. At the LAT and ICP4 loci, wt genomes retained localized interactions, while ΔCTRL2 genomes displayed more distal and dispersed interactions, emphasizing the widespread impact of CTRL2 loss on chromatin looping and regulatory architecture (**S. Fig. 3B&C)**. Together, these results reveal that CTRL2 functions as a critical regulator of both local and long-range chromatin interactions. Its deletion reshapes the viral chromatin landscape, leading to a more accessible and dynamic regulatory environment that may influence HSV-1 transcriptional programs and latency-associated chromatin states.

## Discussion

Virally encoded CTCF insulators are key regulatory elements in the HSV-1 genome, contributing to the maintenance of latency through gene silencing in neurons. Nonetheless, the presence of CTCF-nucleated long-range interactions either in *cis* or *trans* have not been characterized in latent HSV-1 genomes, likely due to experimental limitations associated with existing neuronal latency models. Because large numbers of HSV-1 infected neurons are required for 4C-*seq* applications, neither the *in vivo* models previously used to describe fundamentals of HSV-1 latent biology [57] nor the *in vitro* primary neuronal models generated from mouse ganglia [58] were options for these high-throughput analyses. However, recent advances in establishing HSV-1 quiescence and reliable reactivation in LUHMES cells have revolutionized our ability to perform these complex and high-throughput bioinformatic assays. LUHMES cells have been extensively characterized and used to explore human neurodegenerative diseases [59, 60]. More recently we showed that LUHMES establish HSV-1 quiescence and can be reliably reactivated with the addition of the PI3 kinase inhibitor wortmannin [54]. Further, histone marker composition and miRNA expression are indistinguishable between LUHMES and *in vivo* models of HSV-1 infection, indicating that LUHMES are a reliable and complimentary model to commonly used models of HSV-1 latency, reactivation and chromatin organization [18, 54, 61].

Previous functional characterization of virally encoded CTCF insulators in latent HSV-1 genomes revealed that individual insulators display differential and site-specific insulator activity that includes enhancer-blocking, barrier or silencer function [22, 30]. We previously reported that deletion of the CTRL2 insulator of HSV-1 resulted in the dysregulation of chromatin domains and aberrant (increased) lytic gene expression during latency, suggesting that the insulator was required to maintain IE gene silencing through local control of chromatin compartmentalization around those genome loci [51, 52]. We also showed that the ΔCTRL2 recombinant failed to reactivate *in vivo* and that at 5 days post-reactivation the ΔCTRL2 recombinant displayed significantly attenuated US9 expression [31]. US9 is a leaky-late gene that is required for anterograde axonal transport and reactivation from neurons [62–64] and distance separated from the CTRL2 insulator by over 20kb, and the apparent dependence of the CTRL2 insulator on its expression was intriguing and suggested that the latent HSV-1 genome might be ordered into higher order 3D chromatin structures that would promote efficient gene expression under the right circumstances to allow for reactivation from latency.

Both 4C and UMI-4C-seq methods identified significant differences in the 3D architecture of the ΔCTRL2 recombinant compared to wt genomes, specifically around 3 genomic loci that are known to be required for reactivation (VP16, ICP4 and LAT) [32, 65]. In the absence of the CTRL2 insulator, long-range interactions that map to US8 and US9 are lost and local chromatin domains are disrupted around VP16, ICP4 and the LAT regions suggestive of dynamic regulatory environment that may influence HSV-1 transcriptional programs and latency-associated chromatin states. These findings are consistent with our published data that show that LAT, ICP4 and VP16 gene expression are increased in the absence of the CTRL2 insulator, while the compacted heterochromatin domains that are bounded by the CTRL2 insulator in wt genomes are disrupted in the absence of CTRL2 [51]. Perhaps most intriguing though are the implications that the HSV-1 genome 3D architecture has on the ability of viral genomes to reactivate. We have previously reported that within 2 h of inducing reactivation in latently infected mice, CTCF was differentially evicted from viral genomes in a time frame that precedes the accumulation of lytic transcripts described for reactivation [22, 30]. We also showed using rAAV8 delivery of a CTCF-targeting siRNA that depletion of the CTCF protein in neurons harboring latent HSV-1 resulted in shedding of infectious virus in the absence of other reactivation stressors [66], suggesting that CTCF occupancy was required to maintain genome silencing during latency and that CTCF eviction precedes lytic gene expression and is likely required for efficient reactivation, albeit through unknown mechanisms. Our findings here that show that the deletion of the CTRL2 insulator of HSV-1 results in long-range and broad changes in chromatin looping that reshapes the viral chromatin landscape shows that the 3D structures influence HSV-1 transcriptional programs.

Finally, it is also intriguing to speculate on how these interactions and the 3D organization of wt HSV-1 genomes could promote efficient reactivation. Considering the importance of the CTRL2 insulator in the genome organization of wt virus, we hypothesize that CTCF-nucleated chromatin loops or long-range interactions between the CTRL2 insulator and the previously characterized CTUS1 insulator that is flanking the anterograde transport genes US8 and US9 [21, 31] place the LAT, and specifically, the LAT enhancer elements into close spatial proximity to genes required for anterograde transport. During latency, these insulators are functional enhancer blockers and maintain CTCF binding. However, once reactivation is initiated, CTCF is rapidly evicted, and there is a loss of insulator function so that the LAT enhancer can now activate both IE genes and anterograde transport genes so that the **c**oordinated expression of these genes would ensure that newly-formed viral particles would be produced and transported to corneal epithelial cells **(Fig. 4)**. More extensive mechanistic studies are currently underway to test this model in the context of reactivation *in vivo*.

**Figure 4.**
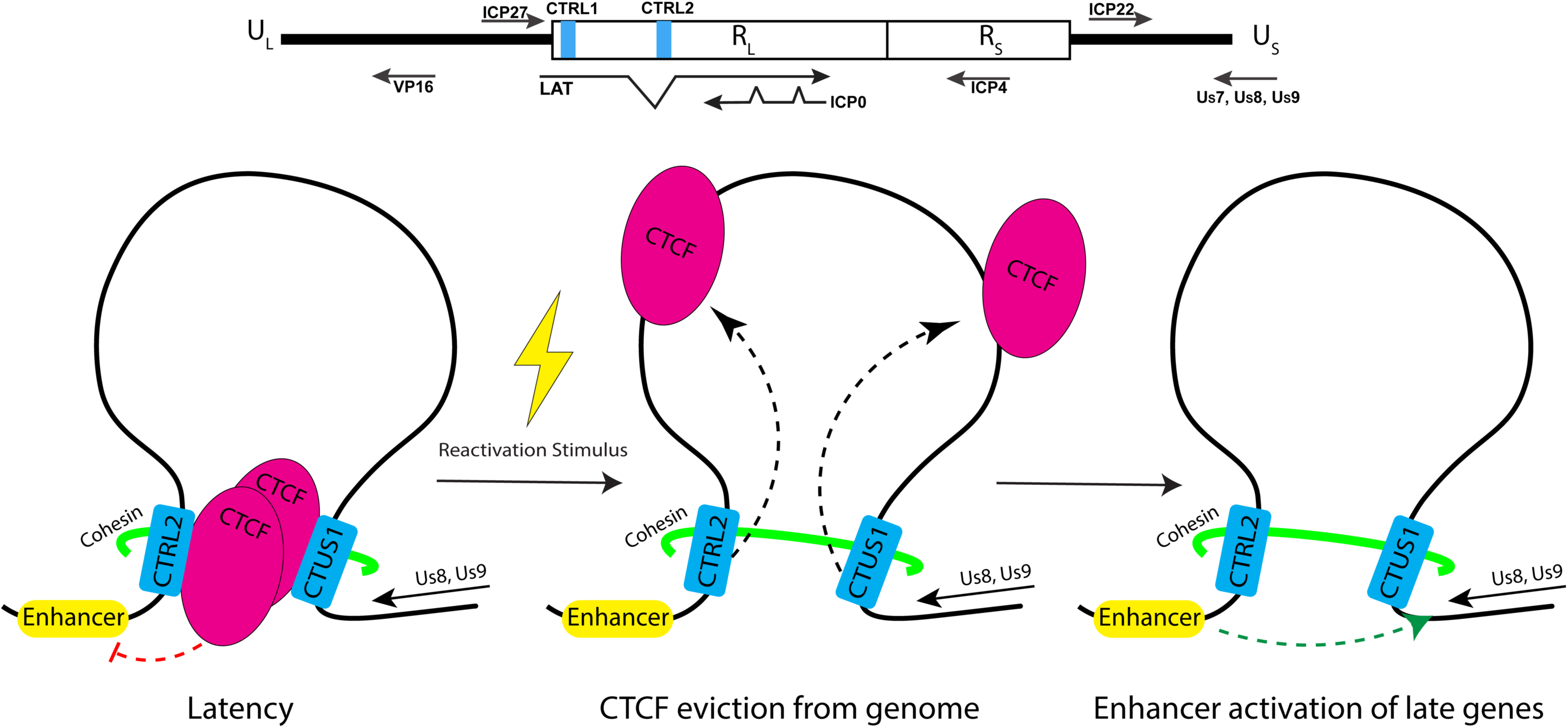
Model for how 3D chromatin architecture contributes to efficient reactivation from latency in HSV-1 infected neurons.

## Materials and Methods

### Cells and Viruses

Lund human mesencephalic (LUHMES) cells were obtained from the ATCC (no. CRL-2927) and were cultured as described previously [54]. All experiments performed in this study utilized LUHMES cells at passages 5 to 7 from the original ATCC stock. Briefly, LUHMES cells were cultured in dishes, on plates, and on glass coverslips, which were coated with poly-L-ornithine hydrobromide (no. P3655; Sigma) overnight at room temperature followed by fibronectin (no. F2006; Sigma) overnight at 37°C. Dishes, plates, and glass coverslips then were rinsed with sterile dH_2_O and allowed to dry overnight. For proliferation, LUHMES cells were propagated in Dulbecco’s modified Eagle’s medium/Ham’s F12 (DMEM/F12; no. 12-719F; Lonza) supplemented with 1× N2 supplement (no. 10378016; Thermo Fisher Scientific), 1% PSG (100 U penicillin, 100 mg/mL streptomycin, 0.292 mg/mL L-glutamine; no. SV30082.01; HyClone), and a 40-ng/mL final concentration of recombinant human fibroblast growth factor (FGF)-basic (no. 100-18B; PeproTech) added fresh to medium before use at 10% CO_2_. Cells were switched to DMEM/F12 supplemented with 1× N2 supplement (no. 10378016; Thermo Fisher Scientific), 1% PSG (100 U penicillin, 100 mg/mL streptomycin, 0.292 mg/mL L-glutamine; no. SV30082.01; HyClone), 1-μg/mL final concentration of tetracycline hydrochloride (no. T7660; Sigma), 1 mM final concentration *N*6,2′-*O*-dibutyryladenosine 3′,5′-cyclic monophosphate sodium salt (no. D0627; Sigma), and a 2-ng/mL final concentration recombinant human glial cell-derived neurotrophic factor (GDNF) (no. 212-GD-010; R&D Systems) at 60% to 70% confluence to induce differentiation as previously described (42). Original stocks of the wild-type HSV-1 strain, 17*Syn+* (GenBank accession number NC_001806), and the recombinant ΔCTRL2 viruses were originally obtained from the Bloom Lab (University of Florida). For the ΔCTRL2 recombinant virus, 135-bp (nt 120,500-120,635) of CTRL2 insulator was deleted from the 17*Syn+* parental wild-type (wt). The recombinant virus was verified by sequence analyses, as previously described [51]. Both virus stocks were propagated on vero cells using a 0.01 MOI. Cultures were supplemented with Eagle’s minimal essential medium with 1% FBS and 1% antibiotic-antimycotic solution at 37°C for 3-4 days and viruses were harvested by centrifugation followed by two freeze thaw cycles. The final supernatants were aliquoted and stored in -80 for further use. To determine viral titers, vero cells were infected in triplicate with 10-fold serial dilutions in DMEM (1% FBS and 1% antibiotic-antimycotic solution) for 72 hrs, plaques were stained by crystal violet and counted. For infections prior to downstream experiments, cells were seeded and grown in 6-well plates to confluency, unless otherwise noted. Monolayers of cells were inoculated with either 17*Syn*+ or the ΔCTRL2 recombinant in DMEM with 1% FBS and 1% antibiotic-antimycotic solution. Plates were rocked at 4 for 1 h to allow for virus adsorption, followed by 30 min incubation at 37, 5% CO_2_. The virus containing media was removed and replaced with fresh DMEM with 1% FBS and 1% antibiotic-antimycotic solution and plates were incubated for the remainder of the indicated time point for cell harvesting.

### LUHMES cells infections

LUHMES cells were grown and differentiated on poly-L-ornithine/fibronectin-coated coverslips (42). Briefly, LUHMES cells were plated at 25,000 cells per well (24-well plates/glass coverslips) or 3 × 10^6^ cells (15-cm dishes) and allowed to proliferate for a period of 2 to 3 days, followed by 5 days of differentiation. To establish a latent infection, the postmitotic neurons were pretreated with 50 μM acyclovir (ACV) (Sigma, PHR1254) for 2 h and infected with HSV-1 viral strain 17*syn*^+^ or ΔCTRL2 recombinant at an MOI of 3 in the presence of 50 μM ACV. After 48 h, medium was removed and replaced with fresh differentiation medium that did not contain ACV. ACV is not included for the remainder of the experiment. Harvesting was performed at the latent time point 8 days post-infection for chromosome conformation capture and downstream sequencing (4C-seq).

### 4C-seq analysis

10 million LUHMES cells, latently infected with HSV-1 strains (wt or ΔCTRL2), were cross-linked using 2% formaldehyde for 10 minutes at room temperature, followed by quenching with 0.125 M glycine and washing with cold PBS. Cell pellets were lysed in a buffer containing Tris-HCl, NaCl, NP-40, Triton X-100, EDTA, PMSF, and protease inhibitors, and homogenized. Nuclei were isolated, pelleted, and resuspended in restriction enzyme buffer and SDS followed by incubation at 37°C. HindIII was used for the first digestion at 37°C for ∼16 hours, and digested samples were inactivated at 65°C and ligated using T4 DNA ligase at 16°C overnight. Digestion and ligation efficiencies was validated with gel electrophoresis. Crosslinks were reversed by adding Proteinase K at 65°C overnight, treatment with RNAseA, followed by phenol-chloroform extraction. Purified DNA was subjected to a second digestion with DpnII and ligated overnight at 16°C to generate circularized DNA fragments. The resulting 4C template was purified using magnetic bead-based cleanup. Inverse PCR was performed using primers designed to flank the restriction enzyme sites and amplify the circularized fragments ligated to the viewpoint. All primers sequences are listed in **S. Table 2**. PCR conditions included 30 cycles with a gradient of annealing temperatures to optimize 4C-seq *cis*-interactions were determined using the program peakC [67]. To visualize interaction differences in chromatin interactions between the two viral genotypes. high sensitivity (HS) assay (Thermo Scientific). Two rounds of nested PCR were performed with bait-specific primers for each viewpoint, yielding libraries of approximately 500 bp. For each library, 5–10 PCR reactions (each with 200 ng of DNA template) were pooled to ensure sufficient complexity. Raw sequencing reads files were demultiplexed and aligned to the HSV-1 reference genome (NC_001806.2) using the UMI4Cats pipeline [68], followed by UMI collapsing to count unique interaction events and remove PCR duplicates. Interaction profiles were normalized to the group with the lowest UMI counts, excluding a 3-kb window surrounding the bait site. Interaction domains were visualized using adaptive smoothing and domainogram representations. Differential interactions were identified using variance stabilizing transformation (DESeq2) [69] and monotone smoothing, with comparisons performed using Wald test and Fisher’s Exact Test for low UMI regions. Differential peaks with a log2 odds ratio >1 were visualized with the Plotgardener R package [70].

## Acknowledgements

This work was supported by the grants NIH/NIAID R01AI134807 (DMN), NIH/NEI 2T32EY027721 (MAS), NIH/NIAID R01AI048633 (DCB), the NSF GRFP 2137424 (KAM), the Core Grant for Vision Research from the NIH to the University of Wisconsin-Madison (P30 EY016665) and an unrestricted grant from Research to Prevent Blindness (Department of Ophthalmology, University of Wisconsin). The funders had no role in study design, data collection and analysis, decision to publish, or preparation of the manuscript.

## Supplemental Figures

**S. Figure 1.** Validation of HSV-1-specific viewpoints for 4C-*seq*. PCR amplification of the bait-targeted viewpoints 1 (VP1) and 2 (VP2) from HSV-1-infected (+) and uninfected (-) 4C templates was performed across three replicates (only first two are shown in the figure). PCR products were resolved on a 1.5% agarose gel, with a DNA ladder shown as a size reference. Amplification was successful in HSV-1-infected samples for both VP1 and VP2, while no amplification was observed in uninfected samples, confirming the specificity of the viewpoints to HSV-1-infected templates.

**S. Figure 3A:** Paired-end sequencing of multiplexed UMI-4C amplicons resulted in a similar number of reads with a relatively balanced distribution of UMIs and a high degree of consistency across replicates across each bait.

**S. Figure 3B:** UMI-normalized interaction frequency profiles across the HSV-1 genome for LAT viewpoints (top panel) and ICP4 (bottom panel) in wt and ΔCTRL2-infected LUHMES cells. Increased interaction frequencies and broader interaction domains are observed in ΔCTRL2 genomes compared to wt.

**S. Figure 3C:** Differential chromatin interaction arcs for ICP4 (top panel) and LAT (bottom panel) viewpoints visualized as arc plots across the HSV-1 genome. Arcs represent statistically significant changes in *cis*-interactions peaks between ΔCTRL2 and wt genomes. Red arcs indicate **increased** interaction frequencies in ΔCTRL2 relative to wt (log odds ratio > 1), while blue arcs indicate **reduced** interactions. HSV-1 genome coordinates and annotated ORFs are displayed below the arc plots.

