## supplemental table 1 for "Multiple long-range *cis* interactions generate CTCF insulator-dependent viral chromatin domains in quiescent HSV-1 genomes"

**Supplementary Table 1: 4C-*seq* peaks coordinates**

| **Sequence** | **Start** | **End** | **Viewpoint** | **Condition** | **Overlap** |
| --- | --- | --- | --- | --- | --- |
| NC_001806.2 | 12582 | 12786 | VP1 | WT | Both |
| NC_001806.2 | 15497 | 15736 | VP1 | WT | Both |
| NC_001806.2 | 27947 | 27969 | VP1 | WT | Both |
| NC_001806.2 | 27971 | 27976 | VP1 | WT | Both |
| NC_001806.2 | 28007 | 28086 | VP1 | WT | Unique |
| NC_001806.2 | 28107 | 28136 | VP1 | WT | Unique |
| NC_001806.2 | 39732 | 39986 | VP1 | WT | Both |
| NC_001806.2 | 80649 | 80836 | VP1 | WT | Both |
| NC_001806.2 | 90047 | 90236 | VP1 | WT | Both |
| NC_001806.2 | 98732 | 98935 | VP1 | WT | Both |
| NC_001806.2 | 133391 | 133536 | VP1 | WT | Unique |
| NC_001806.2 | 138232 | 138486 | VP1 | WT | Both |
| NC_001806.2 | 139941 | 140027 | VP1 | WT | Unique |
| NC_001806.2 | 12591 | 12786 | VP2 | WT | Unique |
| NC_001806.2 | 15497 | 15736 | VP2 | WT | Both |
| NC_001806.2 | 27882 | 28132 | VP2 | WT | Unique |
| NC_001806.2 | 39732 | 39985 | VP2 | WT | Unique |
| NC_001806.2 | 80649 | 80836 | VP2 | WT | Both |
| NC_001806.2 | 90047 | 90236 | VP2 | WT | Both |
| NC_001806.2 | 98732 | 98933 | VP2 | WT | Both |
| NC_001806.2 | 133465 | 133585 | VP2 | WT | Both |
| NC_001806.2 | 138232 | 138386 | VP2 | WT | Unique |
| NC_001806.2 | 138395 | 138486 | VP2 | WT | Unique |
| NC_001806.2 | 143182 | 143286 | VP2 | WT | Unique |
| NC_001806.2 | 12034 | 12184 | VP1 | $\Delta$CTRL2_mutant | Unique |
| NC_001806.2 | 12582 | 12786 | VP1 | $\Delta$CTRL2_mutant | Both |
| NC_001806.2 | 15482 | 15736 | VP1 | $\Delta$CTRL2_mutant | Unique |
| NC_001806.2 | 27947 | 27969 | VP1 | $\Delta$CTRL2_mutant | Both |
| NC_001806.2 | 27971 | 27976 | VP1 | $\Delta$CTRL2_mutant | Both |
| NC_001806.2 | 28007 | 28136 | VP1 | $\Delta$CTRL2_mutant | Unique |
| NC_001806.2 | 39732 | 39986 | VP1 | $\Delta$CTRL2_mutant | Both |
| NC_001806.2 | 80594 | 80836 | VP1 | $\Delta$CTRL2_mutant | Unique |
| NC_001806.2 | 90032 | 90286 | VP1 | $\Delta$CTRL2_mutant | Unique |
| NC_001806.2 | 98732 | 98935 | VP1 | $\Delta$CTRL2_mutant | Both |
| NC_001806.2 | 133382 | 133583 | VP1 | $\Delta$CTRL2_mutant | Unique |
| NC_001806.2 | 138232 | 138486 | VP1 | $\Delta$CTRL2_mutant | Both |
| NC_001806.2 | 138835 | 138936 | VP1 | $\Delta$CTRL2_mutant | Unique |
| NC_001806.2 | 139833 | 140066 | VP1 | $\Delta$CTRL2_mutant | Unique |
| NC_001806.2 | 143182 | 152222 | VP1 | $\Delta$CTRL2_mutant | Unique |
| NC_001806.2 | 12582 | 12786 | VP2 | $\Delta$CTRL2_mutant | Both |
| NC_001806.2 | 15497 | 15736 | VP2 | $\Delta$CTRL2_mutant | Both |
| NC_001806.2 | 27889 | 28123 | VP2 | $\Delta$CTRL2_mutant | Unique |
| NC_001806.2 | 39732 | 39986 | VP2 | $\Delta$CTRL2_mutant | Both |
| NC_001806.2 | 80648 | 80836 | VP2 | $\Delta$CTRL2_mutant | Unique |
| NC_001806.2 | 90040 | 90286 | VP2 | $\Delta$CTRL2_mutant | Unique |
| NC_001806.2 | 98732 | 98933 | VP2 | $\Delta$CTRL2_mutant | Both |
| NC_001806.2 | 133465 | 133585 | VP2 | $\Delta$CTRL2_mutant | Both |
| NC_001806.2 | 138232 | 138436 | VP2 | $\Delta$CTRL2_mutant | Unique |
| NC_001806.2 | 138457 | 138486 | VP2 | $\Delta$CTRL2_mutant | Unique |
