## supplemental table 2 for "Multiple long-range *cis* interactions generate CTCF insulator-dependent viral chromatin domains in quiescent HSV-1 genomes"

**Supplementary Table 2: Primer sequences**

| **Primer Name** | **Sequence** | **Assay** |
| --- | --- | --- |
| VP1_F | TCGTCGGCAGCGTCAGATGTGTATAAGAGACAGATGGTCT  TAACGGCAAGCTT | 4C-seq |
| VP1_R | GTCTCGTGGGCTCGGAGATGTGTATAAGAGACAGTTTTAA  ACACCACCTGCGG | 4C-seq |
| VP2_F | TCGTCGGCAGCGTCAGATGTGTATAAGAGACAGCGGAAC  CCGTGTGCAAGCTT | 4C-seq |
| VP2_R | GTCTCGTGGGCTCGGAGATGTGTATAAGAGACAGCCGCC  CCATCTTAGGAAAA | 4C-seq |
| LAT_US | CAAACACAAACACCCGCGACGG | UMI-4C |
| CTRL2_US | GCAGGCGCTCGCGGAAACTT | UMI-4C |
| VP16_US | CTGCTCAAACTCGAAGTCGGCC | UMI-4C |
| ICP4_US | AGGAACGTCCTCGTCGAGGCGA | UMI-4C |
| LAT_DS | AATGATACGGCGACCACCGAGATCTACACTCTTTCCCTAC  ACGACGCTCTTCCGATCTTCTCCTCGCCTTCTCCCACCCA | UMI-4C |
| CTRL2_DS | AATGATACGGCGACCACCGAGATCTACACTCTTTCCCTACA  CGACGCTCTTCCGATCTCCCAACCCACTGTGGTTCTGGC | UMI-4C |
| VP16_DS | AATGATACGGCGACCACCGAGATCTACACTCTTTCCCTACA  CGACGCTCTTCCGATCTGGGAATCCCCGTCCCCCAACAT | UMI-4C |
| ICP4_DS | AATGATACGGCGACCACCGAGATCTACACTCTTTCCCTAC  ACGACGCTCTTCCGATCTACATTTCCCCAGGCGCTTTTGC | UMI-4C |
| Universal_R | CAAGCAGAAGACGGCATACGA | UMI-4C |
