## supplemental table 3 for "Multiple long-range *cis* interactions generate CTCF insulator-dependent viral chromatin domains in quiescent HSV-1 genomes"

**Supplementary Table 3: UMI-4C peak coordinates**

| **Sequence** | **Start** | **End** | **Width** | **Strand** | **Z-Score** | **P Value** | **Sample** | **Condition** | **Viewpoint** |
| --- | --- | --- | --- | --- | --- | --- | --- | --- | --- |
| NC_001806.2 | 113789 | 116862 | 3074 | * | 1.71472277 | 0.04319803 | $\Delta$CTRL2_latent_rep1_CTRL2 | $\Delta$CTRL2 | CTRL2 |
| NC_001806.2 | 138659 | 142763 | 4105 | * | 1.75441492 | 0.03967972 | $\Delta$CTRL2_latent_rep1_CTRL2 | $\Delta$CTRL2 | CTRL2 |
| NC_001806.2 | 13633 | 17069 | 3437 | * | 1.74750239 | 0.04027512 | $\Delta$CTRL2_latent_rep2_CTRL2 | $\Delta$CTRL2 | CTRL2 |
| NC_001806.2 | 15346 | 19016 | 3671 | * | 1.74750214 | 0.04027514 | $\Delta$CTRL2_latent_rep2_CTRL2 | $\Delta$CTRL2 | CTRL2 |
| NC_001806.2 | 17070 | 21400 | 4331 | * | 1.74750185 | 0.04027516 | $\Delta$CTRL2_latent_rep2_CTRL2 | $\Delta$CTRL2 | CTRL2 |
| NC_001806.2 | 91932 | 94946 | 3015 | * | 2.07741278 | 0.01888174 | $\Delta$CTRL2_latent_rep2_CTRL2 | $\Delta$CTRL2 | CTRL2 |
| NC_001806.2 | 113789 | 116862 | 3074 | * | 2.49433867 | 0.0063096 | $\Delta$CTRL2_latent_rep2_CTRL2 | $\Delta$CTRL2 | CTRL2 |
| NC_001806.2 | 132708 | 135799 | 3092 | * | 2.36749699 | 0.00895443 | $\Delta$CTRL2_latent_rep2_CTRL2 | $\Delta$CTRL2 | CTRL2 |
| NC_001806.2 | 87849 | 89621 | 1773 | * | 1.85034305 | 0.03213206 | WT_latent_rep1_CTRL2 | WT | CTRL2 |
| NC_001806.2 | 110810 | 112910 | 2101 | * | 1.87874399 | 0.03013973 | WT_latent_rep1_CTRL2 | WT | CTRL2 |
| NC_001806.2 | 136225 | 138658 | 2434 | * | 1.77644391 | 0.03782989 | WT_latent_rep1_CTRL2 | WT | CTRL2 |
| NC_001806.2 | 142764 | 144881 | 2118 | * | 1.75239822 | 0.03985268 | WT_latent_rep1_CTRL2 | WT | CTRL2 |
| NC_001806.2 | 13633 | 17069 | 3437 | * | 2.40991491 | 0.00797812 | WT_latent_rep2_CTRL2 | WT | CTRL2 |
| NC_001806.2 | 28796 | 30585 | 1790 | * | 2.34403982 | 0.00953807 | WT_latent_rep2_CTRL2 | WT | CTRL2 |
| NC_001806.2 | 49644 | 51267 | 1624 | * | 1.94380963 | 0.02595921 | WT_latent_rep2_CTRL2 | WT | CTRL2 |
| NC_001806.2 | 107952 | 110809 | 2858 | * | 3.24179957 | 0.00059389 | WT_latent_rep2_CTRL2 | WT | CTRL2 |
| NC_001806.2 | 109522 | 111477 | 1956 | * | 2.3022175 | 0.01066145 | WT_latent_rep2_CTRL2 | WT | CTRL2 |
| NC_001806.2 | 134116 | 136224 | 2109 | * | 1.95055544 | 0.02555498 | WT_latent_rep2_CTRL2 | WT | CTRL2 |
| NC_001806.2 | 135800 | 136718 | 919 | * | 2.76704306 | 0.00282836 | WT_latent_rep2_CTRL2 | WT | CTRL2 |
| NC_001806.2 | 7568 | 11808 | 4241 | * | 1.82541641 | 0.03396911 | $\Delta$CTRL2_latent_rep1_ICP4 | $\Delta$CTRL2 | ICP4 |
| NC_001806.2 | 9488 | 13632 | 4145 | * | 2.1578392 | 0.01547017 | $\Delta$CTRL2_latent_rep1_ICP4 | $\Delta$CTRL2 | ICP4 |
| NC_001806.2 | 52590 | 55240 | 2651 | * | 1.64536861 | 0.04994691 | $\Delta$CTRL2_latent_rep1_ICP4 | $\Delta$CTRL2 | ICP4 |
| NC_001806.2 | 136225 | 138658 | 2434 | * | 2.22565024 | 0.01301881 | $\Delta$CTRL2_latent_rep1_ICP4 | $\Delta$CTRL2 | ICP4 |
| NC_001806.2 | 136719 | 140511 | 3793 | * | 3.22599568 | 0.00062768 | $\Delta$CTRL2_latent_rep1_ICP4 | $\Delta$CTRL2 | ICP4 |
| NC_001806.2 | 138659 | 142763 | 4105 | * | 4.53622216 | 2.8635E-06 | $\Delta$CTRL2_latent_rep1_ICP4 | $\Delta$CTRL2 | ICP4 |
| NC_001806.2 | 140512 | 143353 | 2842 | * | 2.33160195 | 0.00986082 | $\Delta$CTRL2_latent_rep1_ICP4 | $\Delta$CTRL2 | ICP4 |
| NC_001806.2 | 45057 | 49091 | 4035 | * | 2.02390207 | 0.02149011 | $\Delta$CTRL2_latent_rep2_ICP4 | $\Delta$CTRL2 | ICP4 |
| NC_001806.2 | 47857 | 49643 | 1787 | * | 1.83637745 | 0.03315093 | $\Delta$CTRL2_latent_rep2_ICP4 | $\Delta$CTRL2 | ICP4 |
| NC_001806.2 | 113789 | 116862 | 3074 | * | 1.86177061 | 0.03131772 | $\Delta$CTRL2_latent_rep2_ICP4 | $\Delta$CTRL2 | ICP4 |
| NC_001806.2 | 136225 | 138658 | 2434 | * | 1.86239118 | 0.03127399 | $\Delta$CTRL2_latent_rep2_ICP4 | $\Delta$CTRL2 | ICP4 |
| NC_001806.2 | 9488 | 13632 | 4145 | * | 1.97000594 | 0.02441885 | WT_latent_rep1_ICP4 | WT | ICP4 |
| NC_001806.2 | 28178 | 29862 | 1685 | * | 1.78760608 | 0.03691979 | WT_latent_rep1_ICP4 | WT | ICP4 |
| NC_001806.2 | 86980 | 88849 | 1870 | * | 2.51123985 | 0.0060154 | WT_latent_rep1_ICP4 | WT | ICP4 |
| NC_001806.2 | 87849 | 89621 | 1773 | * | 2.15163223 | 0.01571317 | WT_latent_rep1_ICP4 | WT | ICP4 |
| NC_001806.2 | 22985 | 24697 | 1713 | * | 2.44158865 | 0.0073114 | WT_latent_rep2_ICP4 | WT | ICP4 |
| NC_001806.2 | 23657 | 25187 | 1531 | * | 2.43970661 | 0.0073496 | WT_latent_rep2_ICP4 | WT | ICP4 |
| NC_001806.2 | 143354 | 145495 | 2142 | * | 2.05519899 | 0.01992989 | WT_latent_rep2_ICP4 | WT | ICP4 |
| NC_001806.2 | 16792 | 20492 | 3701 | * | 1.76391531 | 0.03887312 | $\Delta$CTRL2_latent_rep1_LAT | $\Delta$CTRL2 | LAT |
| NC_001806.2 | 22637 | 24415 | 1779 | * | 1.68362586 | 0.046127 | $\Delta$CTRL2_latent_rep1_LAT | $\Delta$CTRL2 | LAT |
| NC_001806.2 | 44546 | 48635 | 4090 | * | 1.72770934 | 0.04202017 | $\Delta$CTRL2_latent_rep1_LAT | $\Delta$CTRL2 | LAT |
| NC_001806.2 | 45695 | 49278 | 3584 | * | 1.72975426 | 0.04183709 | $\Delta$CTRL2_latent_rep1_LAT | $\Delta$CTRL2 | LAT |
| NC_001806.2 | 60028 | 64524 | 4497 | * | 2.02294358 | 0.02153948 | $\Delta$CTRL2_latent_rep1_LAT | $\Delta$CTRL2 | LAT |
| NC_001806.2 | 61396 | 65610 | 4215 | * | 1.77258015 | 0.03814915 | $\Delta$CTRL2_latent_rep1_LAT | $\Delta$CTRL2 | LAT |
| NC_001806.2 | 68856 | 73310 | 4455 | * | 2.13116911 | 0.01653761 | $\Delta$CTRL2_latent_rep1_LAT | $\Delta$CTRL2 | LAT |
| NC_001806.2 | 70169 | 75475 | 5307 | * | 1.89308517 | 0.02917327 | $\Delta$CTRL2_latent_rep1_LAT | $\Delta$CTRL2 | LAT |
| NC_001806.2 | 99732 | 103750 | 4019 | * | 2.25861833 | 0.01195357 | $\Delta$CTRL2_latent_rep1_LAT | $\Delta$CTRL2 | LAT |
| NC_001806.2 | 100885 | 104479 | 3595 | * | 2.01736876 | 0.02182852 | $\Delta$CTRL2_latent_rep1_LAT | $\Delta$CTRL2 | LAT |
| NC_001806.2 | 8177 | 12303 | 4127 | * | 2.40371033 | 0.00811481 | $\Delta$CTRL2_latent_rep2_LAT | $\Delta$CTRL2 | LAT |
| NC_001806.2 | 11272 | 14394 | 3123 | * | 2.09366824 | 0.01814477 | $\Delta$CTRL2_latent_rep2_LAT | $\Delta$CTRL2 | LAT |
| NC_001806.2 | 12304 | 16791 | 4488 | * | 2.47572869 | 0.00664823 | $\Delta$CTRL2_latent_rep2_LAT | $\Delta$CTRL2 | LAT |
| NC_001806.2 | 14395 | 18391 | 3997 | * | 2.01050206 | 0.02218904 | $\Delta$CTRL2_latent_rep2_LAT | $\Delta$CTRL2 | LAT |
| NC_001806.2 | 44546 | 48635 | 4090 | * | 1.65748748 | 0.04871048 | $\Delta$CTRL2_latent_rep2_LAT | $\Delta$CTRL2 | LAT |
| NC_001806.2 | 107034 | 110065 | 3032 | * | 1.8552543 | 0.03177996 | $\Delta$CTRL2_latent_rep2_LAT | $\Delta$CTRL2 | LAT |
| NC_001806.2 | 136337 | 139390 | 3054 | * | 1.90102444 | 0.02864941 | $\Delta$CTRL2_latent_rep2_LAT | $\Delta$CTRL2 | LAT |
| NC_001806.2 | 138203 | 141249 | 3047 | * | 3.02549421 | 0.00124114 | $\Delta$CTRL2_latent_rep2_LAT | $\Delta$CTRL2 | LAT |
| NC_001806.2 | 139391 | 142854 | 3464 | * | 2.31275588 | 0.01036803 | $\Delta$CTRL2_latent_rep2_LAT | $\Delta$CTRL2 | LAT |
| NC_001806.2 | 12304 | 16791 | 4488 | * | 1.95562118 | 0.0252549 | WT_latent_rep1_LAT | WT | LAT |
| NC_001806.2 | 21655 | 23226 | 1572 | * | 1.79487676 | 0.03633667 | WT_latent_rep1_LAT | WT | LAT |
| NC_001806.2 | 78540 | 80462 | 1923 | * | 1.70587996 | 0.04401522 | WT_latent_rep1_LAT | WT | LAT |
| NC_001806.2 | 79680 | 81957 | 2278 | * | 1.99601146 | 0.02296634 | WT_latent_rep1_LAT | WT | LAT |
| NC_001806.2 | 100885 | 104479 | 3595 | * | 1.8367193 | 0.03312567 | WT_latent_rep1_LAT | WT | LAT |
| NC_001806.2 | 27685 | 29640 | 1956 | * | 1.69591793 | 0.04495071 | WT_latent_rep2_LAT | WT | LAT |
| NC_001806.2 | 100885 | 104479 | 3595 | * | 2.44540591 | 0.00723446 | WT_latent_rep2_LAT | WT | LAT |
| NC_001806.2 | 103751 | 106328 | 2578 | * | 1.91292179 | 0.02787903 | WT_latent_rep2_LAT | WT | LAT |
| NC_001806.2 | 141250 | 144472 | 3223 | * | 1.87605777 | 0.03032367 | WT_latent_rep2_LAT | WT | LAT |
| NC_001806.2 | 142855 | 145094 | 2240 | * | 2.16168746 | 0.01532114 | WT_latent_rep2_LAT | WT | LAT |
| NC_001806.2 | 91650 | 94380 | 2731 | * | 1.70204701 | 0.04437328 | $\Delta$CTRL2_latent_rep1_VP16 | $\Delta$CTRL2 | VP16 |
| NC_001806.2 | 136415 | 139447 | 3033 | * | 1.84893097 | 0.03223389 | $\Delta$CTRL2_latent_rep1_VP16 | $\Delta$CTRL2 | VP16 |
| NC_001806.2 | 139448 | 143334 | 3887 | * | 1.81690749 | 0.03461564 | $\Delta$CTRL2_latent_rep1_VP16 | $\Delta$CTRL2 | VP16 |
| NC_001806.2 | 8899 | 12506 | 3608 | * | 2.09122794 | 0.01825382 | $\Delta$CTRL2_latent_rep2_VP16 | $\Delta$CTRL2 | VP16 |
| NC_001806.2 | 28054 | 29730 | 1677 | * | 1.859712 | 0.03146314 | $\Delta$CTRL2_latent_rep2_VP16 | $\Delta$CTRL2 | VP16 |
| NC_001806.2 | 115150 | 118193 | 3044 | * | 1.7949743 | 0.0363289 | $\Delta$CTRL2_latent_rep2_VP16 | $\Delta$CTRL2 | VP16 |
| NC_001806.2 | 136415 | 139447 | 3033 | * | 2.83637782 | 0.00228142 | $\Delta$CTRL2_latent_rep2_VP16 | $\Delta$CTRL2 | VP16 |
| NC_001806.2 | 107497 | 110604 | 3108 | * | 1.86651586 | 0.03098461 | WT_latent_rep1_VP16 | WT | VP16 |
| NC_001806.2 | 143335 | 145125 | 1791 | * | 1.69947028 | 0.04461531 | WT_latent_rep1_VP16 | WT | VP16 |
| NC_001806.2 | 8899 | 12506 | 3608 | * | 2.26216429 | 0.01184363 | WT_latent_rep2_VP16 | WT | VP16 |
