## Supplementary figures and images for "Multiple long-range *cis* interactions generate CTCF insulator-dependent viral chromatin domains in quiescent HSV-1 genomes"

### supplemental figure 1

**S. Fig. 1.**

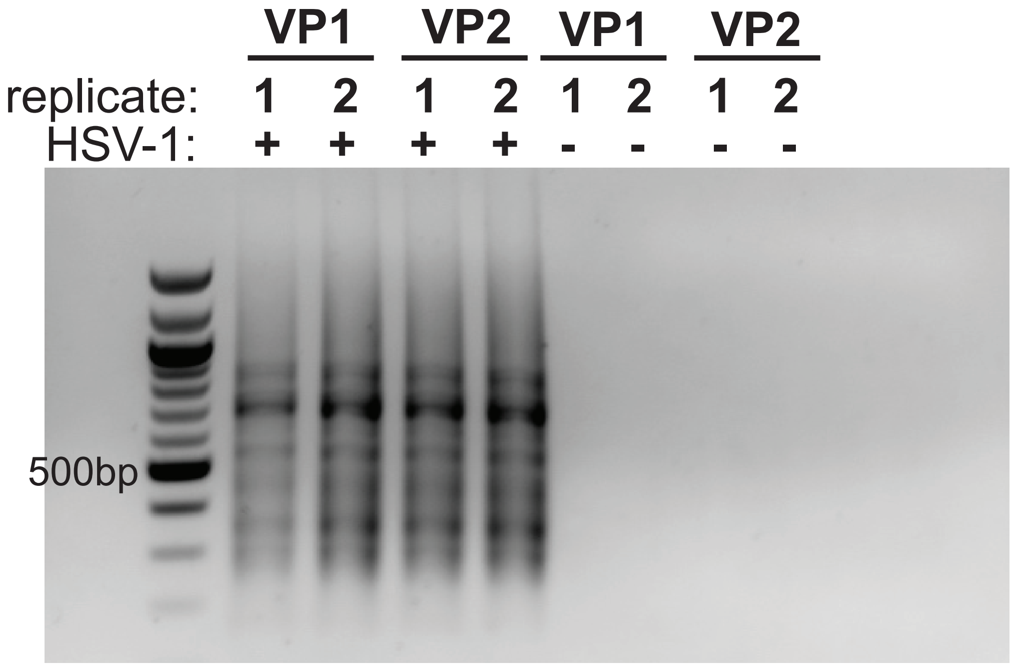

### supplemental figure 3A

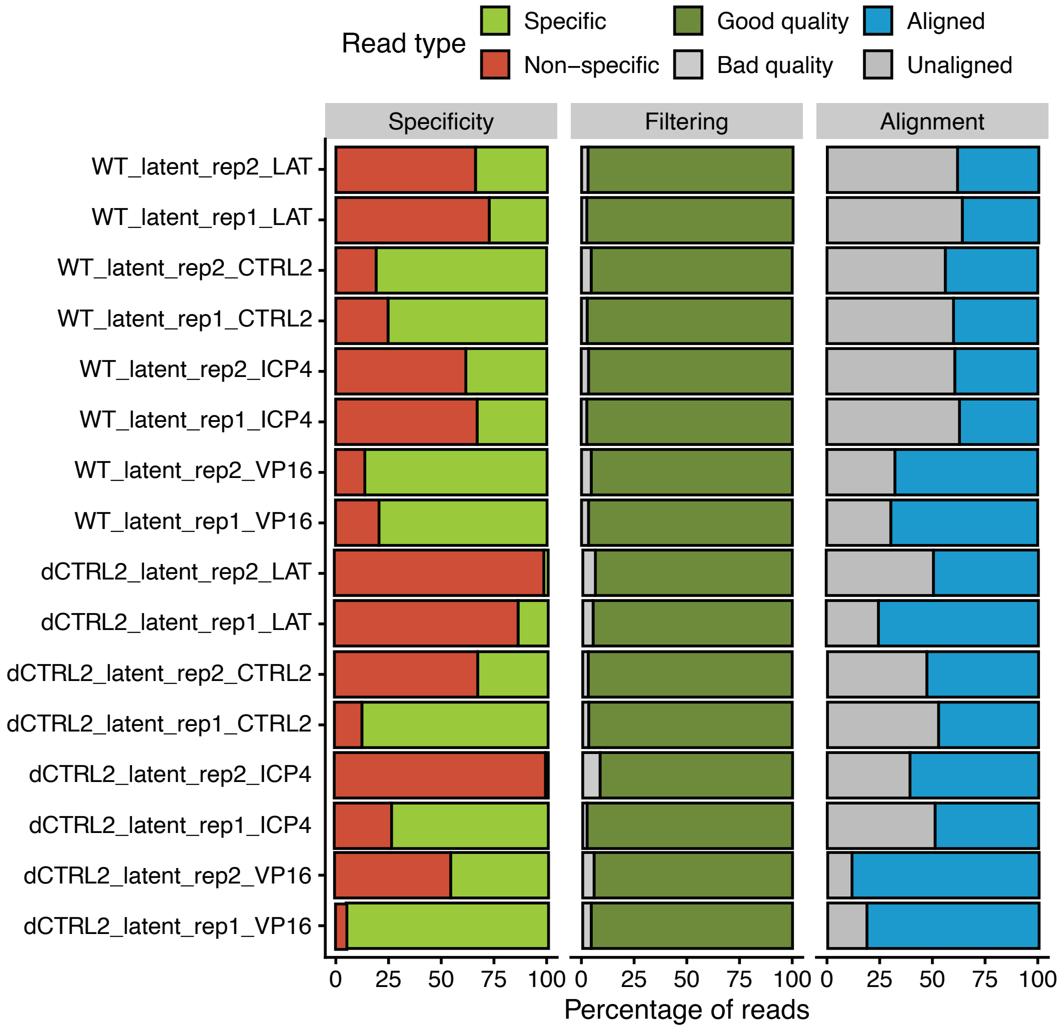

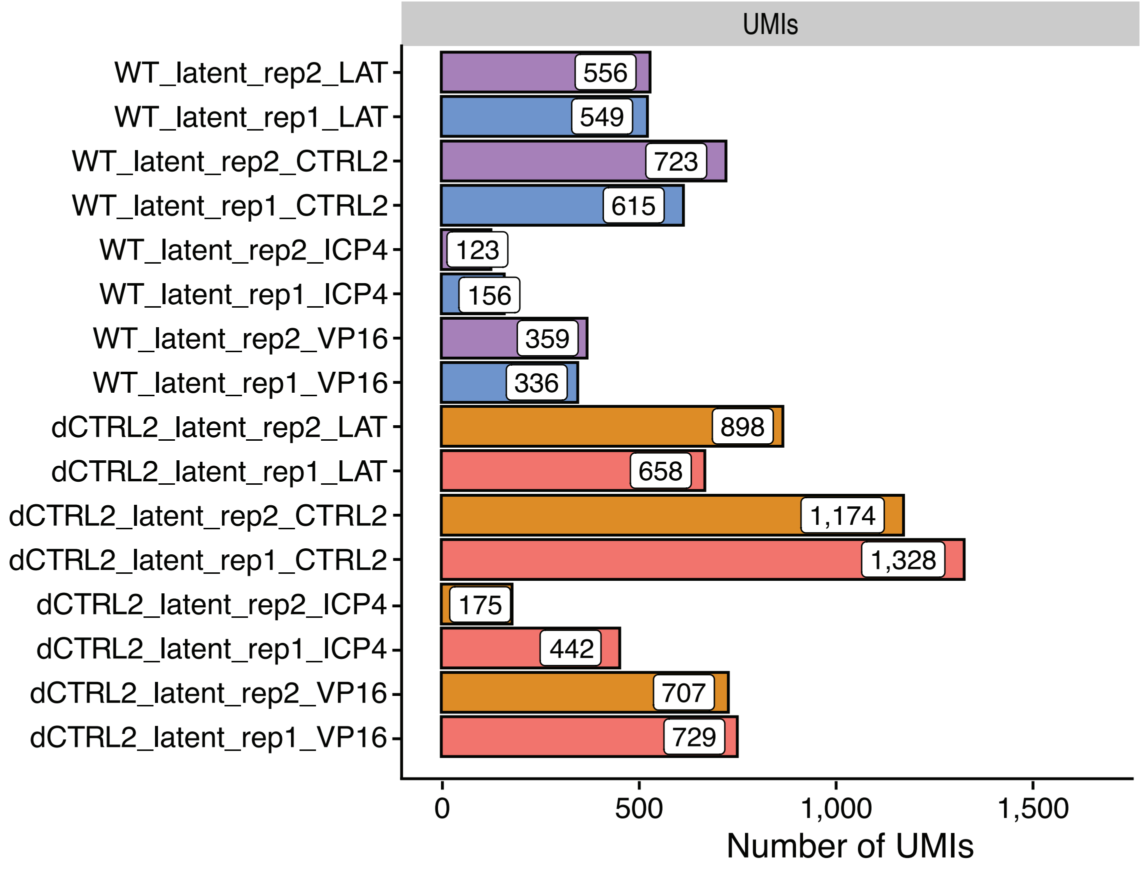

### supplemental figure 3B

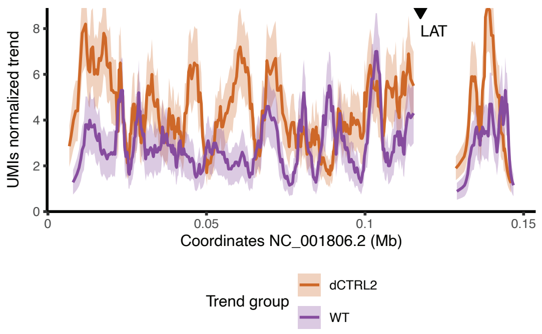

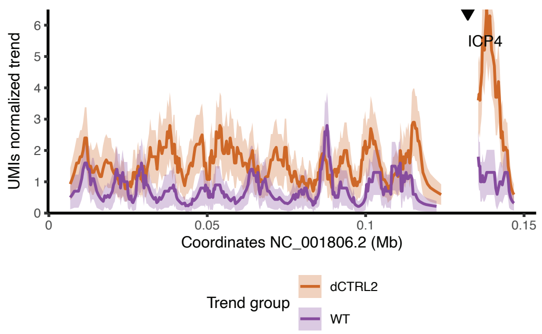

### supplemental figure 3C

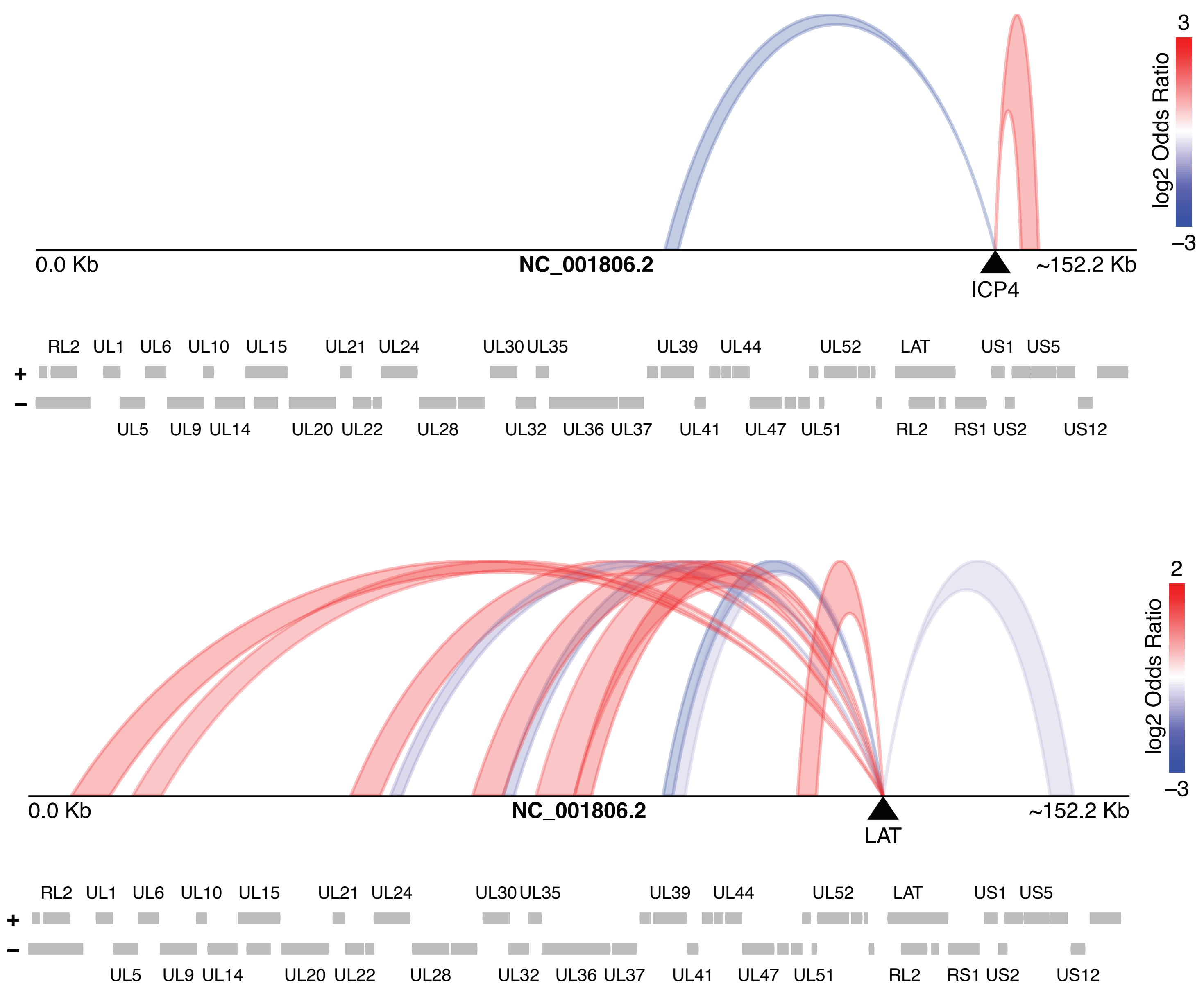
